# Calling genotypes from public RNA-sequencing data enables identification of genetic variants that affect gene-expression levels

**DOI:** 10.1101/007633

**Authors:** Patrick Deelen, Daria V. Zhernakova, Mark de Haan, Marijke van der Sijde, Marc Jan Bonder, Juha Karjalainen, K. Joeri van der Velde, Kristin M. Abbott, Jingyuan Fu, Cisca Wijmenga, Richard J. Sinke, Morris A. Swertz, Lude Franke

## Abstract

Given increasing numbers of RNA-seq samples in the public domain, we studied to what extent expression quantitative trait loci (eQTLs) and allele-specific expression (ASE) can be identified in public RNA-seq data while also deriving the genotypes from the RNA-seq reads. 4,978 human RNA-seq runs, representing many different tissues and cell-types, passed quality control. Even though this data originated from many different laboratories, samples reflecting the same cell-type clustered together, suggesting that technical biases due to different sequencing protocols were limited. We derived genotypes from the RNA-seq reads and imputed non-coding variants. In a joint analysis on 1,262 samples combined, we identified cis-eQTLs effects for 8,034 unique genes. Additionally, we observed strong ASE effects for 34 rare pathogenic variants, corroborating previously observed effects on the corresponding protein levels. Given the exponential growth of the number of publicly available RNA-seq samples, we expect this approach will become relevant for studying tissue-specific effects of rare pathogenic genetic variants.

## Introduction

Most disease-associated genetic variants in humans are regulatory and affect gene-expression levels ^1–3^. With the availability of RNA-sequencing (RNA-seq) two strategies are now commonly used to identify these effects: (1) expression quantitative trait loci (eQTL) mapping to identify common genetic variants that affect gene-expression levels ^4–8^, and (2) allele-specific expression (ASE) analysis to ascertain whether one allele is more abundantly expressed than the other for heterozygous samples. ASE can reveal significant effects even if only a single sample is heterozygous, permitting investigation of rare and low-frequency variants in coding regions. On the other hand, eQTL analyses can be used for any genetic variant, but typically require the use of dozens of samples in order to have sufficient individuals in the different genotype classes ^9–11^. Most eQTL studies so far have focused on a single tissue with large sample sizes ^3,12,13^ (thereby enabling identification of small effects and entire networks of downstream genes, i.e. *trans*-eQTLs) or on a few tissues with limited sample sizes ^14–16^ (enabling identification of tissue- and cell-type-specific *cis-*eQTLs). Although efforts are ongoing, for instance by the GTEx consortium, to investigate larger numbers of different tissues ^17^, still the number of samples studied remains limited. Ideally, eQTL data on many tissues and many different samples should be available, since this would permit eQTL mapping and ASE analyses on rare and low-frequency variants within different cell types. This is especially important for the functional interpretation of clinically important rare variants (particularly recessive Mendelian mutations, where the mutant alleles have appreciable frequencies in the general population ^18^), but will also aid in the classification of variants of unknown significance ^19^.

Fortunately, the raw data of many RNA-seq experiments are being deposited in public databases, and the number of available human RNA-seq samples is growing exponentially, for example, in the European Nucleotide Archive (ENA) (Fig. 1a). Since it has recently been shown that it is possible to derive reliable genotypes from RNA-seq reads ^20^, leveraging publicly available RNA-seq samples might be a viable strategy for obtaining the sample sizes required to perform eQTL mapping and ASE analyses on rare and low-frequency variants across multiple cell-types.

**Figure 1.**
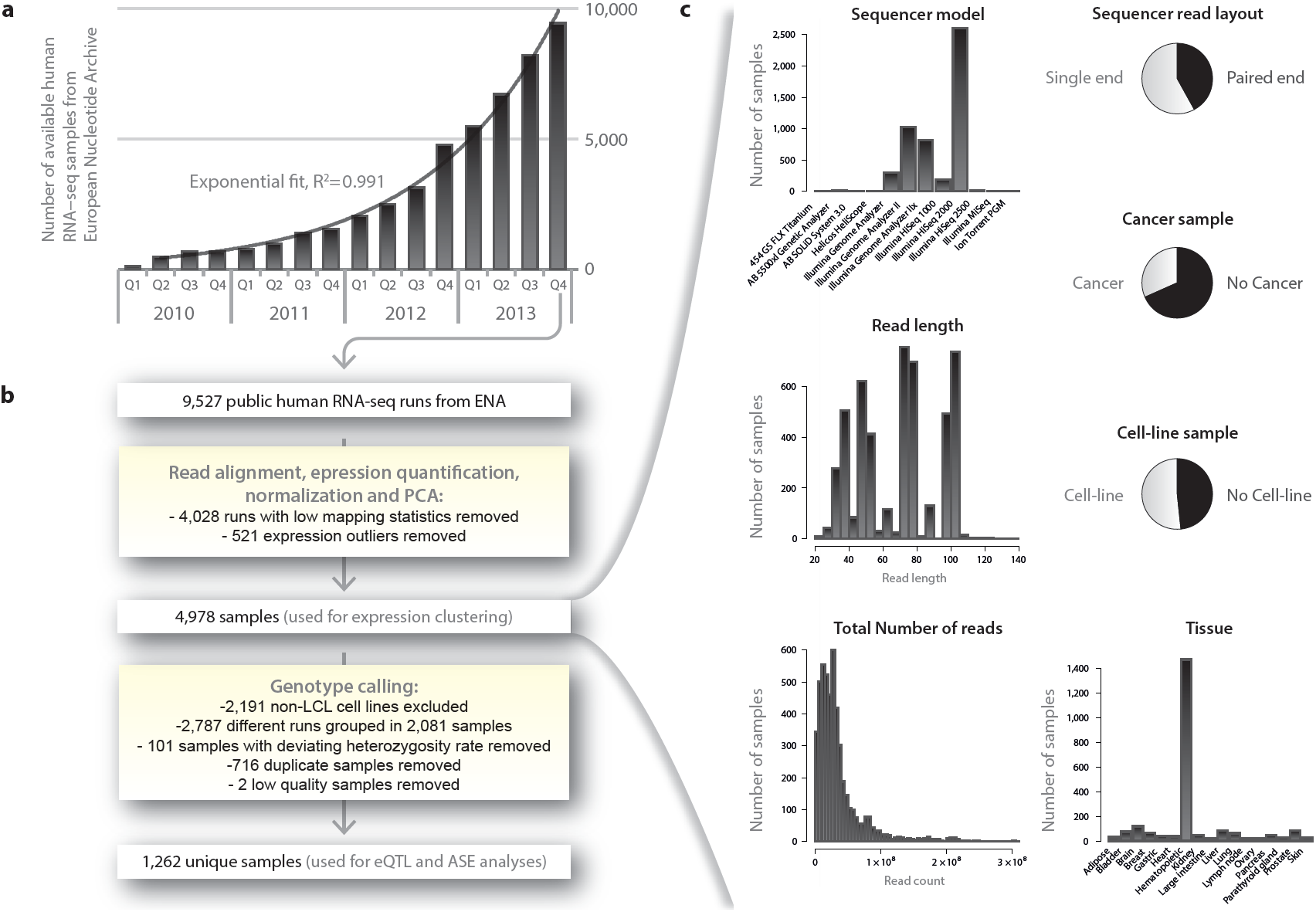
Growth of publicly available RNA-seq and analysis workflow. a) Over the past years the number of available public RNA-seq samples has increased exponentially (exponential fit r^2^ > 0.991) b) General overview of the steps taken to process, quality control and integrate all samples. c) Overview of diversity of 4,978 samples used for expression clustering. Three samples having read lengths >140 (365, 452, 151 bases) are omitted from the read length plot.

Here we present an approach to quantify, normalize and genotype a large number of heterogeneous RNA-seq samples. We show that it is possible to reliably identify eQTLs across many different tissues and also to obtain tissue-specific eQTLs by combining samples from a single tissue derived from many different experiments. We assessed allele-specific expression (ASE) in a large number of samples and identified rare and low-frequency (pathogenic) variants that affect gene-expression levels. We have made all our results freely available online (http://www.molgenis.org/ase), allowing for easy querying of genetic variants of interest.

## Results

### Expression quantification

We downloaded all publicly available human RNA-seq data from the ENA and aligned the reads for each of these samples. We identified 4,978 RNA-seq runs that passed our strict quality control (QC, see Methods). We performed several analyses to ascertain whether these sequence runs, produced in many laboratories around the world, jointly describe biologically coherent patterns. We first conducted principal component analysis (PCA) to obtain a global view on how the different samples clustered together. A PCA on the sample correlation matrix showed that components 1 and 2 permit near-perfect discrimination between primary tissues, cell lines, and hematopoietic tissues (Fig. 2). Other components permit accurate identification of many tissue types such as brain (Fig. 2b, components 4 and 10), liver (Fig. 2c, components 14 and 11) and bladder (Fig. 2d, components 4 and 38), even though the RNA-seq data for these tissues had been generated in at least six different laboratories, with often quite pronounced technical differences (e.g. in sequencer model, read layout, read length, and total number of reads, Fig. 1c). Together, these results indicate that heterogeneous RNA-seq datasets that have been aligned, normalized and QC’ed in a systematic manner yield gene-expression profiles that very clearly describe biologically coherent phenomena. These results also indicate that researchers who would like to learn more about one specific tissue, could combine different (small-scale) RNA-seq data for that tissue into one large dataset.

**Figure 2.**
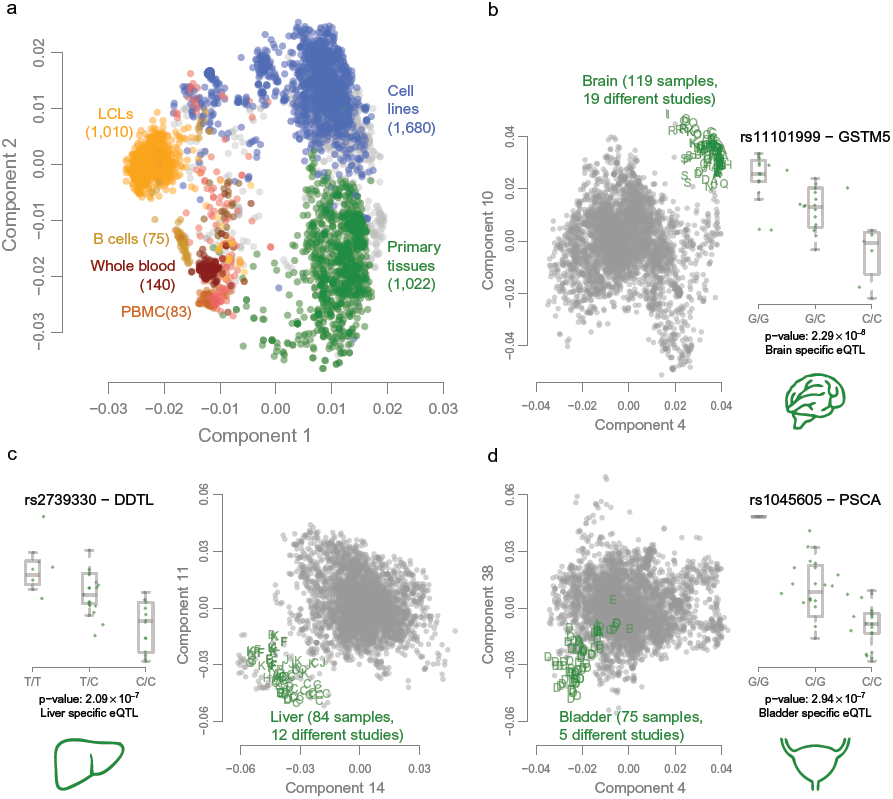
PCA on expression data of all 4,978 samples that passed expression QC. Panel a shows that the first 2 expression components show clear separation between primary tissue samples in green, cell lines (HeLa, K562, Hep G2, etc.) in blue and hematologic tissues and cell types in different shades of red and yellow. The other panels show the two best discriminating components for the different primary tissues. Each letter represents a sample from a distinct study showing that this clustering is not driven by a study specific effect. For each of these three primary tissues, we show an example eQTL effect specific to these tissues.

### Genotyping and imputation

We then assessed whether genotypes could be accurately derived from the samples, which would permit eQTL and ASE analysis. After removing genetically identical samples and additional quality control (see Methods), we had a diverse data set of 1,262 unique individuals (Supplementary Fig. 1). We genotyped 321,415 common SNPs that had a GQ ≥ 30, a call-rate ≥ 80% and a MAF ≥ 0.05. We observed that the total number of high-quality genotype calls that could be made per sample strongly correlated with the total number of sequenced bases per sample (Pearson *r*^2^ = 0.85, Fig. 3a, Supplementary Fig. 2a). As expected, genotypes could only be called in regions where genes are expressed (Fig. 3b, Supplementary Fig. 3).

**Figure 3.**
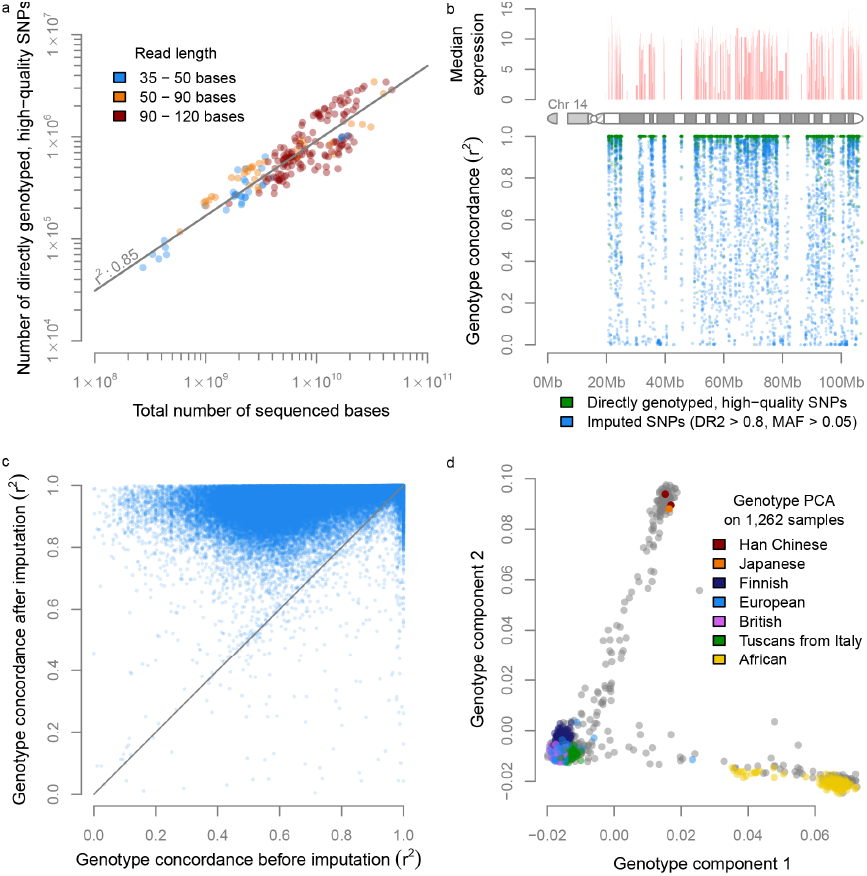
Genotype concordance and genotype PCA on the 1,262 unique individuals that passed genotype QC. a) We observe a strong correlation between the number of sequenced bases and the number of high-quality genotype calls. b) We show that genotyping is only possible in regions with gene-expression c) In 95% of the variants imputation increased genotyping concordance. d) PCA on the genotypes of all 1,262 samples reveals population structure with the expected European, Asian and African clusters.

To ascertain the accuracy of the genotype calls, we compared the RNA-seq-derived genotypes with actual DNA-based genotype calls that were available for 459 Geuvadis ^21^ lymphoblastoid cell-line (LCL) samples that were part of the 1,262 samples. For the ASE analyses we only used high-quality genotype calls (GQ ≥ 30, see Methods), and for this subset of SNPs we observed a median concordance of 1 over all minor allele frequency (MAF) ranges (mean concordance = 0.96).

In order to perform eQTL analysis using non-coding SNPs as well, we used genotype imputation (see Methods) to increase the number of common SNPs to 1,081,155 (predicted dosage *r*^*2*^ (DR2) ≥ 0.8 and MAF ≥ 0.05). Since most of the Geuvadis samples (used for determining the genotyping concordance) are part of the 1000 Genomes Project ^22^, we did not use the 1000 Genome reference panel, but used an independent panel – the Genome of the Netherlands (GoNL) ^23,24^ – to ensure that the genotype concordance measurements were not artificially inflated. The median genotype concordance *r*^*2*^ for the 1,081,155 imputed SNPs for the European Geuvadis samples was *r*^*2*^ = 0.92. When also including the African Geuvadis samples, the genotype concordance decreased somewhat (median *r*^*2*^ = 0.83), because the GoNL imputation reference panel only contained Dutch samples. The genotype concordance of directly genotyped common variants (irrespective of the genotype quality) showed an increased genotype concordance in 95% of the cases after imputation (Fig. 3c). We also observed that, prior to imputation, there is a large difference in genotype concordance of variants in low-expressed genes compared to variants in highly expressed genes, where genotype calling is easier. However, this difference became much smaller after imputation, indicating that it is often possible to accurately call genotypes of SNPs that map within low-expressed genes by using imputation (Supplementary Fig. 4).

PCA on the imputed genotypes confirmed that the major components correctly captured the different ancestries of the individual samples (Fig. 3d). These results also permitted us to stratify the samples into three different groups corresponding to European, African and Asian individuals, and to perform eQTL meta-analyses which are more robust than conducting an eQTL analyses on all samples combined in regions of the genome where allele frequencies differ substantially between populations.

### Cis-eQTL mapping

We then ascertained the reliability of conducting eQTL analysis when using genotypes derived solely from the RNA-seq data. To do so, we tested how many *cis-*eQTLs could be found in the Geuvadis LCL samples when using the RNA-seq derived and imputed genotypes, and also how far they could be replicated using the actual DNA-based genotypes that were available for these samples. An eQTL meta-analysis on the Geuvadis samples using the RNA-seq derived and imputed genotypes (see Methods) resulted in 8,765 unique genes with a significant *cis*-eQTL effect (at a false discovery rate (FDR) ≤ 0.05, Table 1). Of these, 95% could be replicated significantly using the actual DNA-based genotypes (99.95% with the same allelic direction), indicating that eQTL mapping using RNA-seq-derived genotypes is certainly possible for datasets that reflect one sequencing strategy (paired-end 75 bp reads) and one cell type.

**Table 1.**
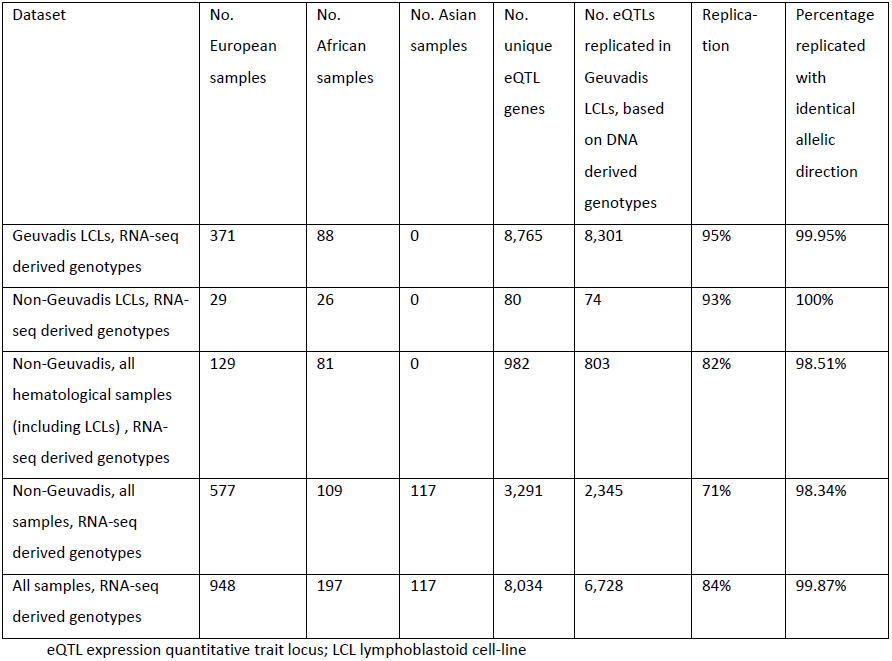
Overview of identified eQTL genes (FDR < 0.05) that were significant in different subsets of the data

| Dataset | No. European samples | No. African samples | No. Asian samples | No. unique eQTL genes | No. eQTLs replicated in Geuvadis LCLs, based on DNA derived genotypes | Replication | Percentage replicated with identical allelic direction |
| --- | --- | --- | --- | --- | --- | --- | --- |
| Geuvadis LCLs, RNA-seq derived genotypes | 371 | 88 | 0 | 8,765 | 8,301 | 95% | 99.95% |
| Non-Geuvadis LCLs, RNA-seq derived genotypes | 29 | 26 | 0 | 80 | 74 | 93% | 100% |
| Non-Geuvadis, all hematological samples (including LCLs) , RNA-seq derived genotypes | 129 | 81 | 0 | 982 | 803 | 82% | 98.51% |
| Non-Geuvadis, all samples, RNA-seq derived genotypes | 577 | 109 | 117 | 3,291 | 2,345 | 71% | 98.34% |
| All samples, RNA-seq derived genotypes | 948 | 197 | 117 | 8,034 | 6,728 | 84% | 99.87% |

eQTL expression quantitative trait locus; LCL lymphoblastoid cell-line

We then performed eQTL analyses (all at FDR ≤ 0.05) on the non-Geuvadis samples while attempting to replicate the identified eQTLs in the Geuvadis data (DNA-based genotypes). We realized that these replication rates would be partly influenced by tissue-specific eQTL effects and first therefore investigated the non-Geuvadis LCL samples (n = 55). Given the sample size, we only identified 80 significant eQTL genes, but we could replicate 93% of these in the Geuvadis samples, all with the same allelic direction (Table 1). Subsequently we performed an eQTL mapping using all the hematological non-Geuvadis samples (n = 210), in which we identified 982 significant eQTL genes, of which 82% could be replicated in the Geuvadis samples (98.51% with identical allelic direction). Finally, we also included the primary tissue non-Geuvadis samples and identified 3,291 significant eQTL genes, of which 71% could be replicated in the Geuvadis LCL samples (98.34% with identical allelic direction).

We then performed an eQTL analysis on all the Geuvadis and non-Geuvadis samples, which identified significant *cis*-eQTLs for 8,034 unique genes (of which 84% were replicated in the Geuvadis DNA-seq-based eQTL data, 99.87% with identical allelic direction). This is fewer than in the analysis on only the Geuvadis samples due to the fact we were dealing with many different tissues in the combined analyses: the small Geuvadis LCL-specific eQTL effects became diluted by the non-LCL samples, leading to fewer *cis*-eQTL effects.

On comparing the 8,765 eQTL genes identified in the Geuvadis samples to the 3,291 eQTL genes identified in the non-Geuvadis samples, we observed that 2,374 genes were identified in both datasets (Fig. 4). As expected, the expression levels of the 903 genes that could not be identified in the Geuvadis data were significantly lower in the Geuvadis data (Wilcox p-value 4.05 × 10^-61^) than in the non-Geuvadis samples.

**Figure 4.**
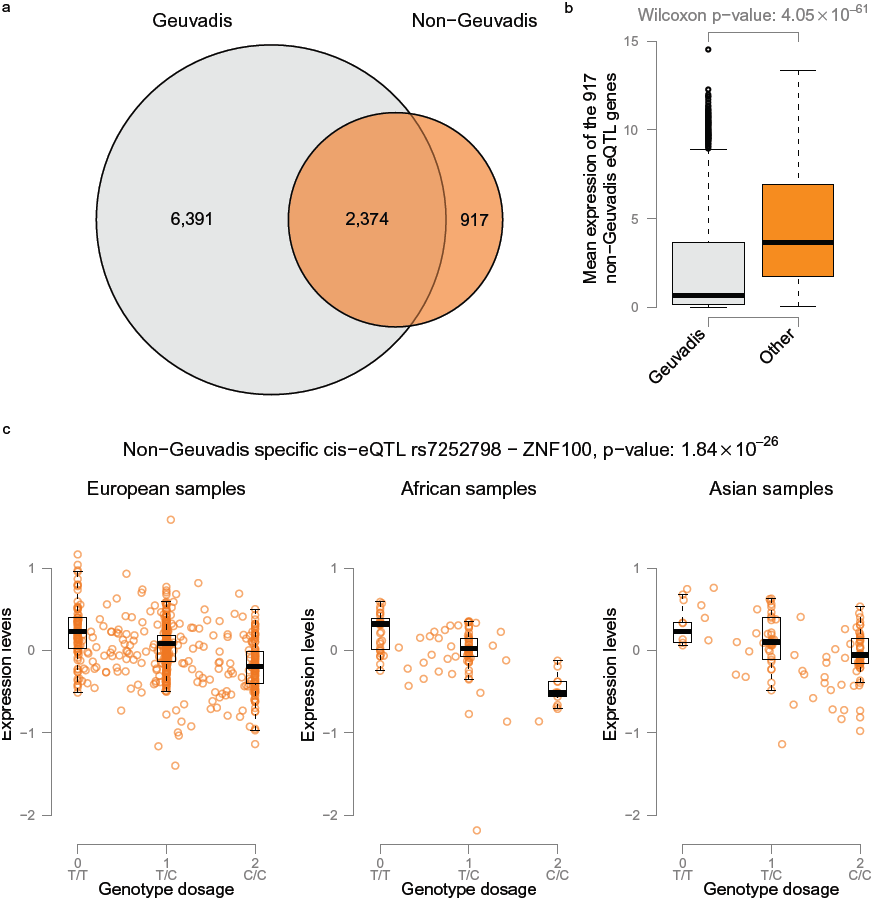
Geuvadis eQTLs vs non-Geuvadis eQTLs. a) When performing eQTL analysis on the non-Geuvadis samples we identify 3,559 significant eQTL genes (FDR < 0.05). 903 of these genes are not detected when using the Geuvadis samples. b) Comparing the expression levels of the genes not identified in the Geuvadis samples we observe that in general these genes are much more abundantly expressed in the non-Geuvadis samples. c) Example eQTL effect of rs7252798 affecting expression levels of ZNF100 that is only identified in the non-Geuvadis samples.

### Tissue-specific eQTL mappings

Since the public RNA-seq data represents many different tissues, we assessed whether it is possible to perform tissue-specific eQTL mapping on samples of the same tissue generated by different laboratories. We performed separate eQTL mapping on four tissues: brain (42 samples from 7 studies), liver (42 samples from 8 studies), bladder (45 samples from 3 studies) and breast (50 samples from 4 studies), since they had eQTL data available on at least 40 samples. This resulted in 121 unique c*is*-regulated genes in brain (32% not detected in Geuvadis), 86 genes in liver (37% not detected in Geuvadis), 65 genes in bladder (38% not detected in Geuvadis) and 43 genes in breast (19% not detected in Geuvadis). As expected, for genes with eQTLs that did not replicate in Geuvadis, we found that the expression in the respective tissues was higher (Supplementary Fig. 5). A representative example is shown for SNP rs11101999, which showed a *cis*-eQTL effect only in brain tissue, on glutathione S-transferase mu 5 (*GSTM5*), a gene that is specifically expressed in brain (Fig. 2b, Supplementary Fig. 6a).

We saw that various GWAS disease-associated genetic variants showed tissue-specific eQTL effects: in the liver samples we found rs2739330 that significantly *cis*-regulates the D-dopachrome tautomerase-like gene (*DDTL*), which is known to be associated with concentrations of liver enzymes in plasma (Fig. 2c, Supplementary Fig. 6b) ^25^. Another example is rs1045605, which affects Prostate stem cell antigen (*PSCA*) gene-expression levels in bladder samples (Fig. 2d, Supplementary Fig. 6c) and which is in near-perfect linkage disequilibrium (LD) with rs2294008 (*r*^*2*^ = 0.98 and *D′* = 0.998), a variant that is associated with both gastric ^26^ and bladder ^27^ cancers.

### Allele*-*specific expression

We mapped ASE by fitting a binomial distribution per SNP using maximum likelihood estimation and then assessed significance by using a likelihood ratio test. Similar to our study of the eQTLs, we first investigated the Geuvadis samples and identified 16,217 ASE SNPs (FDR ≤ 0.05) using the RNA-seq-derived high-quality genotypes (GQ ≥ 30). We compared these results to an ASE analysis using the actual DNA-based genotypes of the Geuvadis samples. 9,221 out of the 9,341 (99%) ASE SNPs that could be tested were replicated using DNA-based genotypes (99.87% with identical allelic direction). Vice versa, on using the Omni DNA genotypes to detect ASE effects, only 232 of them were not found when using RNA-seq genotyping. We next assessed the concordance with the eQTL results: since eQTL mapping and ASE analysis both test the association between genetic variation and gene-expression, we expected the same allele to be more highly expressed in both methods. Indeed, we observed that for 93% of the 1,552 SNPs that showed significant ASE and eQTL effects on the same gene, the allelic direction was consistent. This percentage is similar to another comparison of ASE and eQTL effects, in which 90% of the overlapping eQTL and ASE effects were in the same direction ^28^.

To gain maximum power to detect ASE effects we then performed an analysis on all 1,262 samples and identified 71,214 significant ASE SNPs (FDR ≤ 0.05), of which 4,781 pertained to rare SNPs with a MAF < 0.01 and to 9,018 low-frequency SNPs with a MAF between 0.01 and 0.05. We again compared these ASE SNPs to the eQTL mapping performed on all samples and observed that for 85% of the 1,956 SNPs that showed both significant ASE and eQTL effects on the same gene, the allelic direction was consistent.

It has been reported that nonsense SNPs show ASE with lower expression of the deleterious allele due to nonsense-mediated decay ^9,21^. To investigate this, we annotated the ASE SNPs using SnpEff ^29^ and indeed found that, for nonsense SNPs, the alternative allele is often less expressed than the reference allele (Fig. 5), whereas for other variants we did not observe this bias (Wilcoxon p-value 2.19 × 10^-60^). We also investigated the effect and expected functional impact of the ASE SNPs as predicted by SnpEff. We observed that the SNPs with an expected high functional impact according to SnpEff (Wilcoxon p-value 7.52 × 10^-3^) and those that introduce a stop codon (Wilcoxon p-value 3.66 × 10^-3^) again showed less expression of the alternative allele (Supplementary Fig. 7).

**Figure 5.**
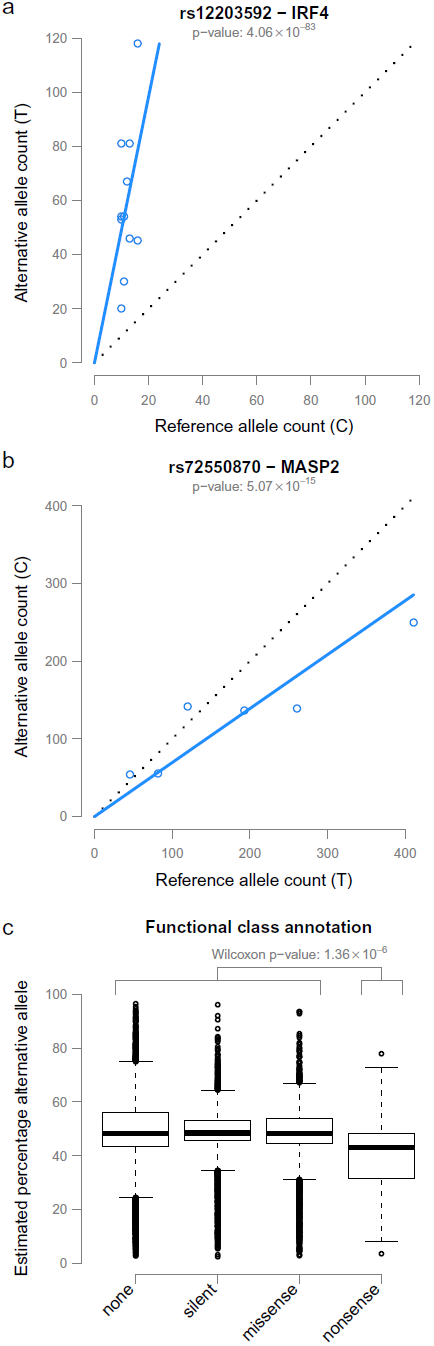
Example ASE effects and direction of ASE effects over different functional classes. a) ASE of rs12203592 located within the *IRF4* gene. The T allele is more abundantly expressed and increases the risk of non-melanoma skin cancers (NMSCs). b) rs72550870, located in *MASP2* shows lower expression for the alternative C allele, known to cause MASP2 deficiency (MASPD). c) All significant ASE SNPs were annotated with functional class information. As expected, nonsense mutations often lead to lower expression levels, in contrast to ASE effects in other functional classes.

We further investigated the functional consequences of common ASE variants by assessing if they were present in the GWAS catalog ^30^. We identified 5 ASE variants with GWAS associations (Supplementary Table 1). For example, rs12203592 is located in the *IRF4* gene and the T allele increases the risk of non-melanoma skin cancers (NMSCs) ^25^ and increases expression levels (Fig. 5a, Supplementary Fig. 8b).

Since ASE mapping also permits the identification of rare and low-frequency variants, we were able to identify 34 variants known to be pathogenic in a Mendelian setting according to the ClinVar database (Supplementary Table 1) ^31^. One example is rs72550870, located in the *MASP2* gene, where we observed an ASE effect (Fig. 5b). It has already been shown that the alternative C allele causes MASP2 deficiency with a recessive inheritance pattern and that heterozygous individuals have significantly lower MASP2 protein levels than individuals homozygous for the wild-type allele ^32^. Our ASE results show exactly the same effect on gene-expression levels. Here it is important to note that the *MASP2* gene is predominantly expressed in the liver (Supplementary Fig. 8a) and that all the samples showing this ASE effect were liver samples, demonstrating the power of a dataset containing multiple tissue types to target a variety of diseases.

The ASE effects obtained can be queried at http://molgenis.org/ase using the MOLGENIS software platform ^33^. It is possible to query a specific variant or search for all ASE variants associated to a specific gene. All our data is available for downloading from this website, including the genotypes, expression data, principal components, eQTLs and ASE effects.

## Discussion

We have shown that it is possible to reliably map eQTLs and perform ASE analyses by calling genotypes directly from RNA-seq data of 1,262 human samples, despite the fact that this data originated from different tissues, was obtained from different laboratories, and was generated using different sequencing techniques.

We called genotypes using GATK, and subsequently imputed using Beagle, while using the GoNL reference panel, to permit unbiased genotype concordance analyses for the Geuvadis samples. We observed that imputation improved the genotype concordance substantially. We find that it is more difficult to impute non-European samples, which is due to our imputation towards the GoNL reference panel. Although GoNL has been shown to yield high-quality imputation for European samples ^23^, its performance has not yet been assessed on non-European populations. It is therefore unreasonable to expect it to perform equally well for Asian or African samples. We expect that a more diverse reference panel will help to resolve this issue in the future.

In this study we had to annotate each sample manually because we found that the annotations available for the different sequence runs were typically limited and inconsistent in terminology. A second reason was that the sample annotations were scattered over multiple databases. Thirdly, although in general the ENA provided better-structured annotations, the information in the Sequence Read Archive was typically more extensive, providing a total of 572 different annotation fields, of which 16 could refer to the tissue of origin. We expect that with more consistency in sample annotation, future large-scale integration of public RNA-seq datasets using automated sample annotation will become feasible.

We have demonstrated that it is possible to run tissue-specific eQTL mapping in public RNA-seq data. We showed that when using only 42 liver samples (originating from eight different labs), it was possible to identify eQTLs that are liver-specific, some of which had been detected by earlier GWAS studies as associated with liver-specific traits. Although the concept of tissue-specific eQTLs is not new, our results demonstrate that different research groups investigating a specific disease in a particular tissue can combine their data in order to conduct joint eQTL mapping. This strategy will certainly prove useful for tissues that are difficult to obtain.

We were able to identify ASE effects for various rare disease-causing variants using only 1,262 samples. We expect our approach will also be useful for studying many other rare pathogenic variants in the near future, because the number of publicly available RNA-seq samples is growing exponentially: at the end of July 2014, the ENA contained 14,831 human RNA-seq samples, which is over 1.5 times the number of samples that we investigated here. Additionally, the read-depth and read-length per sample are steadily increasing (Supplementary Fig. 2b), permitting more sensitive eQTL and ASE analyses (on less expressed genes) on the newly deposited samples.

We anticipate that, with more samples available, eQTL and ASE effects will be detectable for many more (rare) variants. These will be of particular relevance for rare genetic variants that have been identified in patients by exome or genome sequencing but for which the clinical significance remains unknown. If such rare alleles are also present in any of the publicly available RNA-seq samples and they are seen to reduce expression levels strongly, this might suggest they have a loss-of-function effect, strongly warranting clinical follow-up. As such, our approach could well complement existing computational prediction algorithms (that have so far been based primarily on allele frequencies and conservation information), and help speed up the identification of disease-causing mutations, leading to better treatment options and well-informed decisions for patients and their families.

## Methods

### Pipelines and QTL/ASE mapping software

We have made the pipeline and tools that we developed freely available as open source software. The pipelines are implemented in Molgenis compute ^34^ and can be downloaded at: http://github.com/molgenis/molgenis-pipelines. The eQTL/ASE mapping software is publicly available at: http://www.molgenis.org/systemsgenetics/QTL-mapping-pipeline

### Downloading public RNA-seq experiments

We downloaded the samples from the European Nucleotide Archive (ENA). The following filter criteria were used to download the data: Taxon: human (9606), Library strategy: RNA-Seq, Library source: Transcriptomic and Readcount: ≥ 500,000. This was performed on 16 January 2014 and resulted in 9,611 runs. We were able to download FASTQ files for 9,527 runs for which the md5sum was correct after the downloading.

### Read alignment

STAR 2.3.1|^35^ was used to align the reads of the FASTQ files. It is known that read mapping to a common reference creates a mapping bias: more reads will be covering the reference allele than alternative allele ^36^. To correct for this bias and allow the investigation of allelic imbalance, we aligned RNA-seq reads to the reference genome build 37 masked for SNPs with a MAF ≥ 1% in the Genome of The Netherlands (GoNL) data. Only uniquely mapping reads were included. We used a variable number of mismatches per run: for runs with a read length greater than 90 bases we allowed 4 mismatches, for a read length between 60 and 90 we allowed 3 mismatches, and for shorter reads we allowed 2 mismatches. The runs were filtered on their percentage of uniquely mapping reads. We selected 5,499 runs, each having at least 60% uniquely mapping reads and a total of at least 150,000 successfully aligned reads. These filter criteria also ensured that all miRNA experiments were excluded.

### Gene level quantification

We used HTSeq-count 0.5.4 (http://www-huber.embl.de/users/anders/HTSeq/doc/count.html) to quantify gene-expression levels. Ensembl version 71 was used as gene annotation database.

### Identification of gene-expression outliers

We performed quantile normalization and log2 transformation on the data from the 5,499 aligned runs. We then performed a principal component analysis (PCA) over the sample covariance matrix. This revealed 521 strong outliers for the first component (Supplementary Fig. 9). Close inspection of these 521 samples revealed that they included 3 samples that were in fact DNA-seq runs, 312 samples annotated as single-cell sequencing runs, 97 samples that specifically targeted the HLA region and 1 sample was a Geuvadis run^21^ that did not cluster near the other Geuvadis samples. Based on this information we decided to remove these 521 runs leaving 4,978 runs. We then corrected the expression data for GC content and the total number of reads. After standardizing the expression levels for every gene we performed a new PCA (Fig. 2, Supplementary Fig. 10). The raw and normalized expression data and the PCA results can be downloaded here: http://www.molgenis.org/ase. The annotations for each of these runs have been summarized in Supplementary Table 2.

### Genotyping

After removing low quality samples we recalculated the PCA. The first 2 principal components show clear separation between primary tissues, cell lines and hematological tissues (Fig. 2a). To select samples for eQTL and ASE analyses we decided to excluded all tumor-derived cell-line samples (where genotype calling is inherently difficult due to the presence of somatic copy number aberrations), by excluding all non-LCL samples with a principal score > 0 for PC2 (Fig. 2a).

For the genotyping we used a combination of the GATK Unified Genotyper 2.8 ^37^ and imputation of the genotype likelihoods using Beagle 4 r1230 ^38^, which is identical to methods that have been proposed for low-coverage DNA sequencing ^39^. There are many samples that were sequenced using multiple runs, in the cases where this had been specifically mentioned in the sample annotation, we merged all the aligned reads of the different runs to improve genotyping quality. We called genotypes for each sample individually for all 1000 Genomes, GoNL, and ClinVar ^31^ SNPs. We outputted all variants regardless of the calling quality or number of supporting reads. We excluded known RNA-editing sites, variants near splice junctions, and variants at repeat regions, as is recommend when calling variants in RNA-seq data ^20^.

The genotype likelihoods for the variants with a MAF ≥ 1% were used as input for imputation using Beagle 4 with version 5 of GoNL ^23,24^ as a reference. We performed imputation on all the samples merged together. The genotyping concordances of the Geuvadis samples were determined by calculating the correlation between the imputed RNA-seq dosages and the high-quality genotype calls of the Omni2.5 genotyping chips (as generated by the 1000 Genomes project).

For all samples, we calculated heterozygosity rates using the non-imputed genotypes while taking into account only SNPs with MAF ≥ 5% and a read coverage of at least 10 reads. We excluded 100 samples with a heterozygosity rate below 0.2 (suggesting the presence of chromosomal aberrations, uniparental disomies or strong inbreeding) or above 0.4 (suggesting potentially contaminated or pooled samples). This resulted in 1,980 genotyped samples.

### Removing duplicate samples

In order to identify duplicate samples that are not annotated as such by ENA we selected the high-quality imputed genotypes. We selected all variants with an estimated dosage *r*^*2*^ above 0.95, a MAF of 0.05 and a genotyping rate of 0.95. We performed pruning using Plink 1.07 ^40^ (—indep — pairwise 1000 5 0.2) to select independent variants. We then calculated the pairwise genotype concordance for the remaining variants. Based on the resulting distribution we found that a cut-off of 78% was appropriate in order to deem samples duplicates (Supplementary Fig. 11).

If two or more samples were marked as duplicates we gave first priority to the Geuvadis samples, second priority to samples from tissues for which we had most other samples, and finally, those showing the highest number of expressed genes. Among the 1,264 unique samples that were eventually selected there were two samples that we excluded manually because they showed deviating expression levels from what we expected for their presumed tissue and because they had barely passed various other filter criteria. It is also worthwhile to note that these were among the 8 SOLID samples that had passed the rest of the QC. All our criteria finally resulted in 1,262 unique samples that we used for further analyses.

We subsequently investigated *XIST* gene-expression levels and overall chromosome Y expression levels and observed that the expression levels of these samples corresponded well to the gender annotations (available for 41% of the samples, Supplementary Fig. 12).

### Genotype PCA

The genotype PCA was performed on the 1,262 selected unique samples. The variant filtering and pruning was performed using the same settings as for removing the duplicates.

### eQTL mapping

Before we performed eQTL mapping, we selected the best RNA-seq run per sample by choosing the run with the highest number of expressed genes. The runs were normalized using Trimmed Mean of M-values (TMM) ^41^ and log2 transformation, centering and scaling. Finally we corrected our data for the number of mapped reads, the GC percentage, the first 4 genotype components, and the first 100 expression components. We grouped our samples using the genotype PCA into three different groups (Europeans (n = 948), Africans (n = 197) and Asians (n = 117)) and treated this as a meta-analysis when performing the eQTL mapping. We used our previously described eQTL mapping pipeline ^3^ and mapped *cis*-eQTL within 250 kb from the gene center. We only included variants with an expected dosage *r*^*2*^ ≥ 0.8 for the eQTL mapping.

To correct for multiple testing and in order to get reliable false discovery rates, we usually employ a permutation strategy where we define the null-distribution of eQTL effects by randomly assigning the genotype sample identifiers to expression sample identifiers and redoing the eQTL analysis. This is only effective when the genotype data has been generated independently from the expression data, no population stratification exists, and samples reflect the same cell-type or tissue. However, here, the genotype data and expression data have been derived from the same sample, and therefore a different permutation strategy is required, because if a gene is not expressed at all, no genotypes can be derived (and it could well be that subsequent imputation might not be able to resolve this as well either). It is therefore essential to only permute sample identifier labels within sets of samples that reflect the same cell-type or tissue. In order to do so, we permuted the sample identifiers within each of the different studies, because nearly all the studies concentrate on a single tissue. By using this approach we lower the chance that unknown confounders might cause false-positives. This is further alleviated by the fact that we have already accounted for most of the differences in expression between cell-types and tissues by correcting the expression data for 100 principal components.

The replication analysis was performed using the Geuvadis DNA-seq samples where we treated each Geuvadis population separately in a meta-analysis. For each replication analysis we only tested the most significant SNP for each significant gene.

We also performed tissue-specific eQTL mapping in four tissues. We selected only the samples coming from one population, which resulted in 42 European brain samples, 50 European breast samples, 42 European liver samples, and 45 Asian bladder samples. We ran eQTL mapping in the same manner as described above, with the exception that we performed a normal permutation since all samples were from the same tissue, and we tested whether the identified eQTLs were detectable in the Geuvadis LCL eQTL data.

### Allele-Specific Expression analysis

We performed allele-specific expression (ASE) analysis by fitting per SNP a binomial distribution using maximum likelihood estimation and subsequently assessed significance by using a likelihood ratio test. The FDR was controlled using the Benjamini–Hochberg procedure. During our initial ASE analysis (not shown) we observed a strong reference bias for low-frequency variants that had not been previously masked. We therefore again performed the masking of the reference genome using all 1000G, GoNL and ClinVar variants and performed a new mapping of the 1,262 samples selected for the ASE analysis. We used Samtools mpileup 0.1.19 ^42^ and a custom script to obtain the read counts from these bam files, using only bases with a quality score of at least 17. We excluded variants in known RNA-editing sites, near splice junctions and in repeats regions in the same way as when we did the genotyping.

We checked for each sample genotype if the GATK deemed the individual heterozygous for this variant, thereby only using genotypes with a phred-scaled genotype quality (GQ) score above 30. For ASE analysis we selected the SNPs that were heterozygous in at least 5 samples, had at least 10 reads per allele, and at least 2% of all reads supporting each allele. We removed the sites that had a mappability score < 1 according to the USCS mappability track (CGR Alignability of 50mers) ^21,43^. Using these criteria to select variants we tested for ASE in 56,825 SNPs when only interrogating the Geuvadis samples and tested for ASE in 225,562 SNPs when using all 1,262 samples.

## List of abbreviations

eQTL: Expression quantitative trait locus
ASE: Allele-specific expression
ENA: European nucleotide archive
MAF: Minor allele frequency
RNA-seq: RNA-sequencing
PCA: Principal component analysis
QC: Quality control
LCL: Lymphoblastoid cell-line
FDR: False discovery rate
GoNL: Genome of the Netherlands
GQ: Phred-scaled genotype quality
DR2: Estimated dosage *r*^*2*^ after imputation

## Authors’ contributions

PD and DVZ performed most of the data processing and analysis. PD, DVZ and LF designed the study and wrote the manuscript. LF and MAS supervised the work. MdH, MvdS, MJB, JK, JF and JvdV performed part of the analyses. KMA, CW and RJS contributed to the study design and discussion.

## Acknowledgements

We thank the Target project (http://www.rug.nl/target) for providing the compute infrastructure, the MOLGENIS developers Fleur D. L. Kelpin and Bart Charbon for help with the online query tool, Jackie Senior for editing the manuscript, and Ye Chun for helpful comments. This work was supported by the Netherlands Organization for Scientific Research [NWO-VENI grant 916.10.135 to LF and NWO-VIDI grant 016.146.374 to LF], and a Horizon Breakthrough grant from the Netherlands Genomics Initiative [grant 92519031 to LF]. The research leading to these results also received funding from the European Community’s Health Seventh Framework Programme (FP7/2007–2013) under grant agreement no. 259867. This study was financed in part by the SIA-raakPRO subsidy for the BioCOMP project, and in part by Rainbow grants 2 and 3 from BBMRI-NL, a research infrastructure financed by the Netherlands Organization for Scientific Research [NWO project 184.021.007 to LF, MS, PD, DVZ].

